# Structural determinants underlie the essentiality of the *m*-type thioredoxin TrxA in *Synechocystis*

**DOI:** 10.64898/2026.11.05.764636

**Authors:** Manuel J Mallén-Ponce, Jesús Moron-López, Francisco J Florencio, María José Huertas

## Abstract

Thioredoxins are central regulators of thiol-based redox signaling in photosynthetic organisms, where multiple isoforms with partially overlapping functions coexist. Among them, TrxA is the cyanobacterial homolog of chloroplast *m*-type thioredoxins and the only thioredoxin known to be essential in cyanobacteria. However, it is unclear whether this essentiality is due to its high intracellular abundance or to its specific molecular properties. Here, we evaluated the ability of different thioredoxins to functionally replace TrxA and used TrxA/TrxB chimeras to identify the structural determinants underlying its functional specificity. Overexpression of TrxB, TrxQ or TrxC did not support complete segregation of a disrupted *trxA* locus, although the proteins accumulated to levels comparable to those of endogenous TrxA. Similarly, heterologous expression of NTRC did not rescue the phenotype associated with reduced TrxA levels. Comparative structural analysis revealed differences in surface electrostatics and structural features surrounding the catalytic site of TrxA compared with the other thioredoxins. Analysis of TrxB/TrxA chimeras showed that determinants required for functional complementation are primarily located within the C-terminal region of TrxA. Chimeric variants also differed in their ability to activate cyanobacterial FBP/SBPase and pea chloroplast FBPase *in vitro*, supporting a role for this region in target-specific interactions. Together, these results demonstrate that the essential function of TrxA cannot be explained by protein abundance alone and requires molecular determinants located primarily within its C-terminal region.

## INTRODUCTION

Thioredoxins (Trxs) are small soluble oxidoreductases widely distributed across all domains of life (Schürmann and Buchanan, 2008). Characterized by a conserved Trx fold and a catalytic active site containing the canonical WC(G/P)PC motif, Trxs play a central role in cellular redox homeostasis by catalyzing dithiol-disulfide exchange reactions that regulate numerous cellular processes (Holmgren et al., 2005). In oxygenic photosynthetic organisms, Trxs have diversified into several phylogenetic and functional types. Three major lineages are conserved between cyanobacteria and photosynthetic eukaryotes, including the cyanobacterial TrxA, TrxB and TrxQ, which are related to chloroplast *m*-, *x*- and *y*-type Trxs, respectively (Mallén-Ponce et al., 2022). TrxA is the only Trx present in all cyanobacteria and photosynthetic eukaryotes, which also contain multiple TrxA or *m*-type Trx isoforms. In cyanobacteria, the duplication of TrxA in many lineages and the high similarity of these TrxA paralogues suggest that these functions were originally fulfilled by a single copy. Its duplication may subsequently serve as a backup of essential functions.

In oxygenic photosynthetic organisms, Trxs are mainly reduced by the ferredoxin-thioredoxin reductase (FTR) system, which links photosynthetic electron transport to thiol-dependent redox regulation. FTR is a heterodimeric enzyme composed of a catalytic subunit containing a [4Fe-4S] cluster (FTRc) and a variable subunit (FTRv), with ferredoxin serving as the electron donor (Balsera et al., 2013). FTR is widely distributed among cyanobacterial lineages and is also present in chloroplasts, although it is absent in some cyanobacteria, such as *Prochlorococcus* and *Gloeobacter*, where alternative ferredoxin-dependent reductases have been identified (Buey et al., 2017b; Mallén-Ponce et al., 2022).

Flavin-thioredoxin reductases constitute another group of Trx reductases and include NAD(P)H-thioredoxin reductases (NTRs), which use NADH or NADPH as electron donors (Balsera and Buchanan, 2019). Cyanobacteria also contains ferredoxin flavin-thioredoxin reductase (FFTR), an NTR-related enzyme that lacks the NADPH-binding site and is reduced by ferredoxin. FFTR is the only identified Trx reductase in *Prochlorococcus* and *Gloeobacter* (Buey et al., 2017a). Many cyanobacteria and chloroplasts also contain NADPH thioredoxin reductase C (NTRC), a bifunctional enzyme comprising an NTR module fused to a Trx domain (Serrato et al., 2004). NTRC does not appear to interact with Trxs, but it efficiently reduces 2-Cys peroxiredoxin (2-Cys Prx) in both cyanobacteria and chloroplasts (Pérez-Ruiz et al., 2006; Pascual et al., 2011).

Trx-based redox regulation controls key cellular processes including carbon assimilation, antioxidant response and protein synthesis. Early biochemical studies in photosynthetic organisms identified several Calvin-Benson (CB) cycle enzymes as Trx targets, including phosphoribulokinase (PRK), glyceraldehyde 3-phosphate dehydrogenase (GAPDH) or fructose-1,6-bisphosphatase (FBPase) (Michelet et al., 2013; Mallén-Ponce et al., 2022). In cyanobacteria, redox regulation of carbon metabolism involves both conserved and lineage-specific targets. PRK retains redox regulation across the green lineage, while cyanobacteria also possess a bifunctional fructose-1,6-bisphosphatase/sedoheptulose-1,7-bisphosphatase (FBP/SBPase) enzyme that is regulated by TrxA (Gerbling et al., 1986; Mallén-Ponce et al., 2021). CP12, a conserved small intrinsically disordered protein, is also subject to Trx-dependent redox regulation and forms an inhibitory complex with PRK and GAPDH in its oxidized state (Wedel et al., 1997; Gurrieri et al., 2021). Proteome-wide measurements in *Synechocystis* further indicate that CB enzymes operate at distinct redox states *in vivo*; consistent with a hierarchical redox network shaped by cellular context and and Trx interactions (Tanaka et al., 2026).

In addition to CB cycle regulation, TrxA also links photosynthetic redox control to the oxidative pentose phosphate pathway by indirectly inactivating glucose-6-phosphate dehydrogenase (G6PDH) through reduction of its allosteric activator OpcA. Structural and redox-proteomic studies have shown that OpcA regulates G6PDH through allosteric conformational changes and redox-sensitive cysteine modifications near the protein-protein interface (Doello et al., 2024). Together, these findings illustrate the broad role of TrxA in coordinating redox control of central metabolic pathways in cyanobacteria, extending beyond the classical regulation of CB cycle enzymes.

Despite the high conservation of the Trx fold and catalytic motif throughout evolution, several studies have provided evidence for functional specificity among Trx types. In chloroplasts, redox analyses of some CB cycle enzymes revealed a higher affinity for *f*-type Trxs than for *m*-type Trxs (Wolosiuk et al., 1979; Michelet et al., 2013; Yoshida et al., 2015). Other Trxs, such as *x*- and *y*-type Trxs, appear to regulate the oxidative stress responses (Collin et al., 2003; Collin et al., 2004; Vieira Dos Santos et al., 2007). Recent protein-engineering work in *Chlamydomonas* further supports the idea that Trx specificity can be altered by modifying selectivity determinants outside the catalytic cysteines, highlighting the importance of surface features and target-recognition regions in defining Trx function (Lemaire et al., 2025). In cyanobacteria, previous studies have also suggested functional differentiation among Trx isoforms, although this aspect remains largely unexplored (Pérez-Pérez et al., 2009). However, most reports have been primarily based on *in vitro* biochemical studies, where even different Trxs can reduce the same target protein. Subsequent *in vivo* studies, including the generation of knockout or knockdown mutant strains for one or two Trxs, suggested a high degree of overlap between the target proteins of the different Trxs. Indeed, most Trx mutants in plants and cyanobacteria display phenotypes only under specific conditions (Geigenberger et al., 2017; Mallén-Ponce et al., 2022). In contrast, TrxA (*m*-type Trx) mutants exhibit phenotypes even under standard growth conditions (Wang et al., 2013; Okegawa and Motohashi, 2015; Mallén-Ponce et al., 2021).

Many studies have not considered that TrxA (*m*-type Trx) is the most abundant Trx in photosynthetic organisms. Analyses of Trx levels in *Synechocystis* and *Anabaena* revealed that TrxA represents about 80% of the total Trx pool (Florencio et al., 2006; Mihara et al., 2020), whereas in *Arabidopsis* it constitutes approximately 70% of the total Trx pool (Okegawa and Motohashi, 2015). These observations, together with the inability to obtain mutant strains completely lacking this Trx (Muller and Buchanan, 1989; Navarro and Florencio, 1996), suggested that TrxA essentiality could derive mainly from its high intracellular abundance. Recent work with a conditional TrxA mutant has shown that reduced TrxA levels impair protein synthesis and alter carbon/nitrogen partitioning, reinforcing the idea that TrxA controls central cellular processes beyond individual metabolic enzymes (Mallén-Ponce et al., 2024).

Together, these observations raise the question of whether TrxA essentiality depends primarily on its abundance or on intrinsic molecular properties that determine target recognition. Here, we show that TrxA essentiality cannot be explained by intracellular abundance alone. Instead, genetic, structural and biochemical analyses of TrxA/TrxB chimeras reveal that conserved surface regions downstream of the active site are required for functional interaction with redox regulated targets.

## MATERIALS AND METHODS

### Strains and culture conditions

*Synechocystis* sp. PCC 6803 was grown at 30 °C in BG11 supplemented with 12 mM NaHCO₃ (BG11C) (Rippka R, 1988), bubbled with 1% (v/v) CO₂ in air under continuous illumination (50 μmol photons m⁻² s⁻¹). Sodium arsenite (NaAsO₂; 1 mM) was added when required to induce the *arsB* promoter.

For inducer removal experiments, exponentially growing cells cultured with 1 mM arsenite were harvested (4,200 × g, 5 min), washed twice with BG11C and resuspended at a similar cell concentration in fresh medium with or without inducer. Solid BG11C contains 1% (w/v) agar. Antibiotics were used at the following concentrations: nourseothricin and kanamycin, 50 μg mL⁻¹; spectinomycin and streptomycin, 2.5 μg mL⁻¹ each.

### Generation of mutant *Synechocystis* strains

The *trxA* (*slr0623*), *trxB* (*slr1139*), *trxQ* (*slr0233*), and *trxC* (*sll1057*) genes were PCR-amplified, and chimeric TrxA/TrxB constructs were generated by overlap-extension PCR using the oligonucleotides listed in Table S3. After sequence verification, chimeric constructs were cloned into pglnN::ParsB via *EcoR*I*/Not*I restriction sites; native *trx* genes were cloned directly into the same vector. These constructs target the *glnN* (*slr0288*) locus and confer nourseothricin resistance (Tables S2 and S5).

The constructs were introduced into WT *Synechocystis* and, where indicated, the SR background. To assess TrxA essentiality, strains were subsequently transformed with pGQ5.4+ (Mallén-Ponce et al., 2021), which disrupts the endogenous *trxA* gene. Fully segregated strains were obtained for STXA3, STXBA and STXB48A.

For NTRC expression, the *ntrc* gene (*all0737*) from *Anabaena* sp. PCC 7120 was excised from p28AnNTR2 using *Nde*I*/Not*I and cloned into pnrsD::PcpcB, generating pBS-NTRC. This construct targets the *nrsD* (*slr0796*) locus and confers kanamycin resistance. Transformation of WT and STXA2 generated strains SNTRC and STXA2C, respectively.

### PCR verification and chromosome segregation analysis

Genomic DNA from *Synechocystis* transformants was used to verify construct integration and chromosome segregation by PCR. Integration of the *arsB*-driven constructs at *glnN* and the *PcpcB-ntrC* construct at *nrsD* was assessed using primers flanking the respective insertion sites. Segregation of the endogenous *trxA* locus was assessed using primers distinguishing the WT and disrupted alleles. PCR products were separated by agarose gel electrophoresis and visualized by DNA staining. All primers are listed in Table S3.

### Protein extraction and immunoblot analysis

10 mL cultures were harvested and frozen at -20°C. Cell pellets were resuspended in 50 mM Tris–HCl, pH 8, 25 mM NaCl and 1 mM Phenylmethylsulfonyl fluoride. Subsequently, cells were lysed using glass beads and total proteins were extracted as previously described (Reyes et al., 1995). Total protein concentration was determined using the Lowry assay (Markwell et al., 1978). 15 µg of proteins were separated by sodium dodecyl sulfate polyacrylamide gel electrophoresis (SDS-PAGE), transferred to nitrocellulose membranes (Bio-Rad) and immunoblotted with the indicated antibodies in each case (Table S4). Finally, blots were analyzed using the ECL Prime Western Blot Detection Reagent (Bio-Rad) with secondary anti-rabbit antibody (1:20.000) following the instructions of the manufacturer.

### Cloning, expression and purification of recombinant chimeric Trxs

Recombinant TrxA and TrxB were purified as previously described (Navarro and Florencio, 1996; Pérez-Pérez et al., 2009). Chimeric Trxs and the *Synechocystis* FBP/SBPase (García-Cañas et al., 2022) and pea FBPase (Pla and Lopez-Gorge, 1981) target enzymes were produced in *E. coli* BL21(DE3) using the corresponding pET28-derived plasmids described above and in Table S5. Protein expression was induced with 0.2 mM IPTG, followed by incubation for 24 h at 30 °C.

Cells were harvested, resuspended in 50 mM Tris–HCl (pH 8.0), 150 mM NaCl, 5 mM imidazole and 1 mM PMSF, and disrupted by sonication. His-tagged proteins were purified by Ni-NTA affinity chromatography, eluted with the same buffer containing 500 mM imidazole and desalted using PD-10 columns equilibrated with 50 mM Tris–HCl (pH 8.0) and 150 mM NaCl. Proteins recovered predominantly in the insoluble fraction were purified under denaturing conditions and refolded using a urea-based protocol.

Proteins were further purified by size-exclusion chromatography using an ENrich SEC70 column connected to a Bio-Rad NGC system. Fractions containing the target proteins were analyzed by SDS–PAGE, pooled and stored at −20 °C in 10% (v/v) glycerol. Protein concentrations were determined by absorbance at 280 nm using a NanoDrop spectrophotometer and theoretical extinction coefficients calculated with ProtParam. When required, proteins were concentrated using Amicon Ultra centrifugal filters with a 10 kDa molecular-weight cutoff.

### Fructose-1,6-bisphosphatase activity assay

Purified *Synechocystis* FBP/SBPase and pea FBPase were oxidized by overnight incubation at 4 °C with continuous agitation before the assays. Enzyme activity was determined by measuring the inorganic phosphate released from fructose-1,6-bisphosphate using the Fiske– Subbarow method (García-Cañas et al., 2022). The reagent contained 9 N H₂SO₄, 0.053 M (NH₄)₆Mo₇O₂₄·4H₂O and 0.30 M FeSO₄·7H₂O. Absorbance was measured at 660 nm, and phosphate concentrations were calculated using an inorganic phosphate standard curve.

Reaction mixtures contained 1 µM FBP/SBPase or pea FBPase, 1 µM recombinant Trx or chimeric Trx, 15 mM MgSO₄·7H₂O, 15 mM Tris–HCl (pH 8.0) and 50 µM DTT. After preincubation for 15 min at room temperature, reactions were initiated by adding fructose-1,6-bisphosphate to 7.15 mM and incubated at 30 °C for 8 min. Reactions were stopped with 1 mL Fiske–Subbarow reagent, and absorbance was measured at 660 nm.

### Structural predictions and computational processing

Structures of Trxs and Trx–target complexes were predicted using AlphaFold 3. The four highest-ranked models for each prediction were evaluated based on the predicted template modeling (pTM) score, interface pTM (ipTM) score for complexes, and predicted aligned error (PAE). Models with pTM and ipTM scores above 0.7 and favorable PAE plots were selected for analysis.

Structures were visualized using UCSF ChimeraX. Electrostatic surface potentials were calculated using APBS after structure preparation with PDB2PQR (Jurrus et al., 2018), and protein–protein interfaces were analyzed using PICKCLUSTER (Genz et al., 2025).

Protein net charges at pH 7.5 were calculated using Protein Calculator, and theoretical isoelectric points were obtained using ExPASy ProtParam.

### Statistical analysis

Data from at least three biological replicates were analyzed using GraphPad Prism. Enzymatic activities obtained with each Trx or chimeric Trx were compared with the DTT-only control (No Trx) by one-way ANOVA followed by Dunnett’s multiple-comparisons test.

## RESULTS

### TrxA possesses specific functions beyond its protein abundance

Several proteomic analyses have revealed a high degree of overlap among the target proteins of different Trxs. However, despite this apparent functional redundancy, multiple attempts to eliminate TrxA have been unsuccessful. One possible explanation is that TrxA is more abundant than other Trxs (Mallén-Ponce et al., 2022). To assess whether TrxA essentiality results from this higher cellular abundance, we overexpressed different Trxs and evaluated their ability to compensate for TrxA function.

TrxA, TrxB, TrxQ and TrxC were expressed under the control of the arsenite-inducible *arsB* promoter, generating the parental overexpression strains STXA3P, STXBOEP, STXQOEP and STXCOE1P, respectively (Fig. 1a-b). Unlike the previously characterized STXA2 strain (Mallén-Ponce et al., 2021), these constructs retain the original ribosome-binding site upstream of the *arsB* promoter, while the endogenous *trxA* gene remains intact (Fig. S1). In addition to the wild type (WT) background, the same constructs were introduced into the SARSR (SR) background, which lacks the *arsR* gene encoding the repressor of the *arsBHC* operon (López-Maury et al., 2003). This generated the SRTXA3P, SRTXBOEP, SRTXQOEP and SRTXCOE1P parental strains (Fig. S2a). In contrast to WT-derived overexpression strains, SR-derived strains exhibit constitutive expression of the *arsBHC* operon and do not respond to arsenite (hereafter inducer).

**Figure 1.**
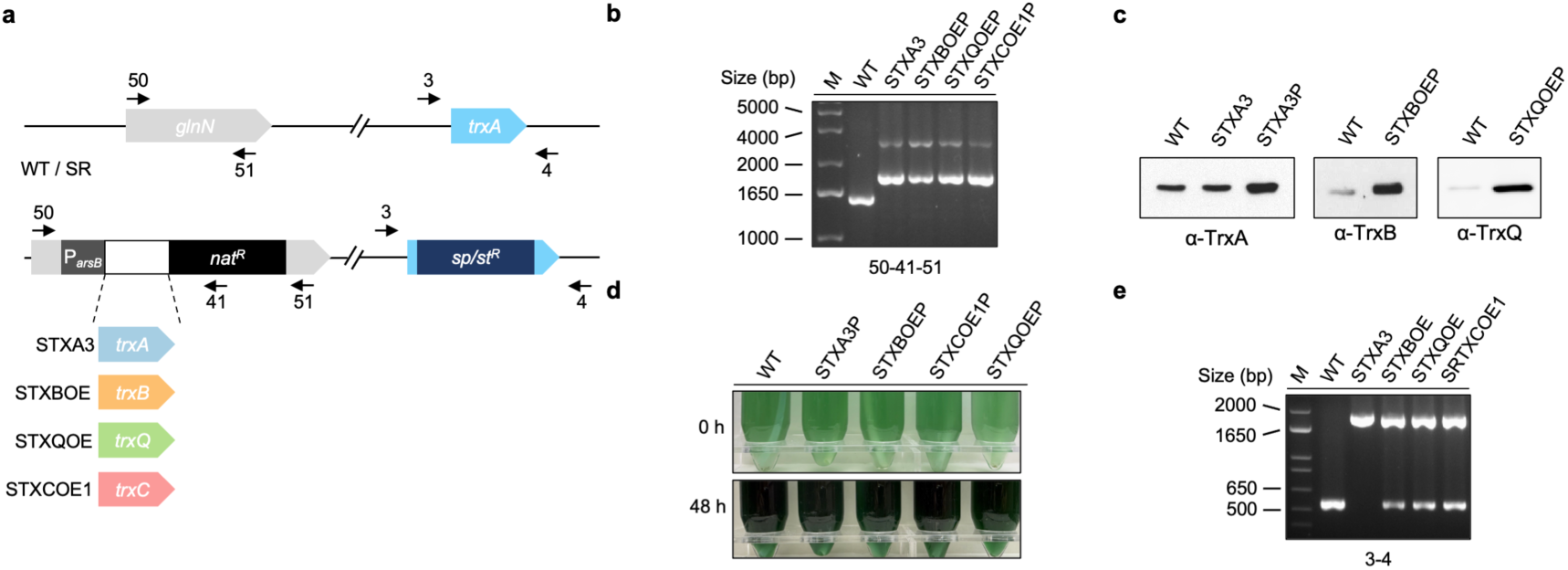
Construction and characterization of *Synechocystis* strains overexpressing thioredoxins. **(a)** Schematic representation of the genetic constructs used for arsenite-inducible expression of TrxA, TrxB, TrxQ and TrxC in the WT and SR backgrounds. **(b)** PCR analysis of the parental strains used to generate the corresponding trxA-disrupted derivatives. The numbers indicate the primers used. **(c)** Western blot analysis of TrxA, TrxB and TrxQ in the WT, STXA3P and STXA3 strains. **(d)** Phenotype of WT, STXA3P, STXBOEP, STXCOE1P and STXQOEP strains at 0h and 48 h in the presence of inducer. **(e)** PCR analysis of *trxA* segregation in the indicated strains. M, molecular size marker. Strains carrying the disrupted *trxA* gene are named without the “P” suffix. Alt text: Multi-panel figure showing schematic diagrams of the *Synechocystis* strains overexpressing thioredoxins TrxA, TrxB, TrxQ and TrxC, agarose gels of PCR products verifying the strains and *trx*A segregation, a western blot detecting TrxA, TrxB and TrxQ, and photographs of liquid cultures of the different strains at 0 and 48 h after induction.

Western blot analysis confirmed that TrxA protein levels in the parental STXA3P strain were approximately 2-fold higher than in the WT strain (Fig. 1c). In the parental STXBOEP and STXQOEP strains, TrxB and TrxQ protein levels were approximately 7-and 35-fold higher than in the WT strain, respectively. These increases resulted in TrxB and TrxQ protein levels comparable to those of endogenous TrxA in WT cells, consistent with the expression levels reported for these Trxs in previous studies (Florencio et al., 2006; Baers et al., 2019; Wang et al., 2022; Spät et al., 2023). Comparable protein levels were observed in the SRTXA3P, SRTXBOEP and SRTXQOEP strains, where TrxA expression occurs constitutively (Fig. S2b). Despite Trx overexpression, no detectable phenotypic alterations were observed (Fig. 1d, Fig. S2c).

The endogenous *trxA* gene was then disrupted in these parental backgrounds by insertion of a spectinomycin/streptomycin resistance cassette. PCR analysis showed that the disrupted *trxA* gene could not be fully segregated in the STXBOE, STXQOE and STXCOE1 strains or in their corresponding SR-derived strains (Fig. 1e, Fig. S2d), indicating that TrxB, TrxQ and TrxC cannot functionally replace TrxA even when expressed at protein levels comparable to those of TrxA. Several attempts to achieve complete chromosome segregation were unsuccessful, even at 1 mM inducer, a concentration sufficient to achieve maximal expression from the *arsB* promoter. In contrast, complete segregation of the disrupted *trxA* locus was achieved in the STXA3 and SRTXA3 strains (Figs. 1e, Fig. S2d), which accumulated TrxA at levels comparable to those in the WT and SR, respectively (Fig. 1c, Fig. S2b).

These TrxA levels were sufficient to prevent any detectable phenotype in the STXA3 strain upon inducer removal (Fig. S2e-f), unlike the previously characterized STXA2 strain (Mallén-Ponce et al., 2022). Given that the inability of TrxB, TrxQ and TrxC to replace TrxA appears to reflect specific functions of TrxA rather than differences in protein abundance, we asked whether an alternative, structurally distinct thiol-reducing system could be bypass this requirement. Although NTRC is absent in *Synechocystis*, this enzyme is present in other cyanobacteria and chloroplasts, where it contributes to thiol-based redox regulation using NADPH as the source of reducing power. Its physiological role in cyanobacteria and chloroplasts has been the subject of intense debate in recent years (Cejudo et al., 2021; Mallén-Ponce et al., 2022). Because NTRC contains both Trx reductase and Trx domains within a single polypeptide, we investigated whether its expression in *Synechocystis* could compensate for reduced TrxA levels or functionally replace the FTR/TrxA pathway. To this end, strains expressing NTRC from *Anabaena* sp. PCC7120 were generated in the *Synechocystis* WT and STXA2 backgrounds, yielding the SNTRC and STXA2C strains, respectively. For overexpression, the *ntrC* gene was placed under the control of the constitutive *cpcB* promoter (Fig. S3a), and complete segregation of NTRC and STXA2C strains was confirmed by PCR analysis (Fig. S3b). Subsequently, expression of NTRC protein was confirmed by western blot analysis (Fig. S3c). Growth analysis revealed that the NTRC expression did not increase the growth rate, and no phenotypic changes were observed (Fig. S3d-e). Likewise, no differences in growth were observed between the STXA2C and STXA2 strains, either in the presence of the inducer or after its removal. These data indicate that heterologous NTRC does not compensate for reduced TrxA levels or functionally replace FTR/TrxA system under these conditions.

### Surface properties distinguish the different types of thioredoxin

To better understand the structural basis underlying functional specificity of the different Trxs, we conducted a comparative analysis of their sequence and surface-related properties. All native Trxs contain a fairly similar number of residues at the N- and C-terminal sides relative to the active site (Table 1, Fig. S4). In contrast, their net charges vary among different Trxs (Table 1). TrxQ has a net charge of 0, while TrxA, TrxB, and TrxC have net charges of -5.1, -7.1, and - 5.2, respectively. Accordingly, TrxA and TrxB have similar theoretical pI values, whereas TrxC and particularly TrxQ differ from this pattern (Table 1). The unusually high pI of TrxQ likely reflects the presence of multiple basic residues. Pairwise sequence identities among the four Trxs are relatively low, ranging from 21.5% to 39.1%, with TrxC showing the lowest overall identity to the other *Synechocystis* Trxs (Table S1, Fig. S4).

**Table 1.** Amino acid distribution, net charge and theoretical pI of *Synechocystis* thioredoxins and chimeric variants.

| **Protein** | **N-side residues** | **C-side residues** | **Net charge** | **Theoretical pI** |
|---|---|---|---|---|
| **Native Trxs** |  |  |  |  |
| TrxA | 31 | 72 | -5.1 | 4.69 |
| TrxB | 30 | 75 | -7.1 | 4.90 |
| TrxQ | 29 | 72 | 0.0 | 7.99 |
| TrxC | 28 | 78 | -5.2 | 5.88 |
| **Trx Chimeric variants** |  |  |  |  |
| TrxAB | 31 | 75 | -8.1 | 4.70 |
| TrxBA | 30 | 72 | -4.1 | 4.98 |
| TrxB48A | 30 | 72 | -2.2 | 6.27 |
| TrxB63A | 30 | 72 | -2.2 | 6.25 |
| TrxB86A | 30 | 72 | -4.0 | 5.34 |

*Note: N-side and C-side indicate the number of amino acid residues located before and after the active site, respectively.*

To further explore structural determinants of specificity, we analyzed the surface properties of individual Trxs. The structures of *Synechocystis* Trxs were predicted using AlphaFold (Jumper et al., 2021; Evans et al., 2022). Consistent with previous studies (Eklund et al., 1991), all Trxs adopt the conserved Trx fold consisting of a central β-sheet flanked by α-helices (Fig. 2), with the catalytic motif exposed on their surface. Moreover, all Trxs display a mixed polarity around the catalytic cysteines, including TrxC and its unconventional active site motif (WCGLC). In TrxA, the acidic residues Asp61 and Glu62 form a mildly electronegative patch on α-helix 3 that is highly conserved among photosynthetic organisms (Mallén-Ponce et al., 2022). In addition, a conserved loop comprising residues 70-76 contains a conserved Arg73 whose guanidine group protrudes from the molecular surface (Fig. 2, Fig. S4). These structural features suggest that surface-exposed residues may therefore play an important role in target recognition.

**Figure 2.**
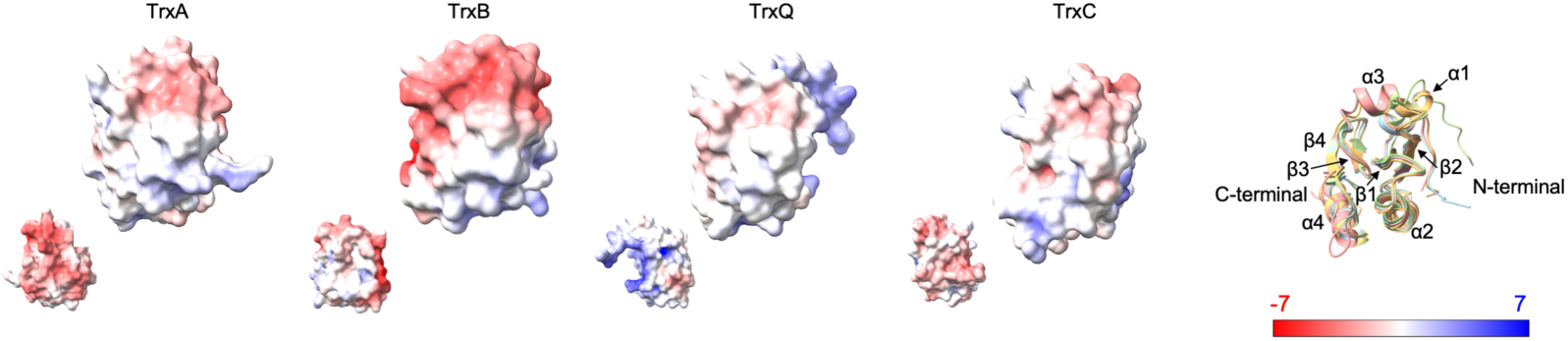
Electrostatic surface properties of *Synechocystis* thioredoxins. Surface representations of the AlphaFold-predicted structures of TrxA, TrxB, TrxQ and TrxC. Electrostatic potentials were calculated using the APBS algorithm and are shown from electropositive (blue) to electronegative (red). Smaller representations show the opposite face of each Trx. The conserved Trx fold and the orientation of the catalytic site are maintained in all proteins, whereas each Trx displays distinct surface electrostatic properties. The right panel shows a representative ribbon structure indicating secondary structure elements (α1-α4 and β1-β4), N- and C-terminal ends. Alt text: Molecular surface models of *Synechocystis* thioredoxins (TrxA, TrxB, TrxQ and TrxC) colored by electrostatic potential from blue (positive) to red (negative), each shown from two opposite faces, with all four sharing the same fold but differing in surface charge distribution. A ribbon diagram on the right labels the helices α1–α4, strands β1–β4 and the N- and C-terminus.

The surface of TrxB is more similar to that of TrxA, although TrxB exhibits a larger electronegative potential on the α-helix 3 and β-strand 3 side of the active site owing to the presence of several acidic residues, including Glu58, Glu72 and Glu90 (Fig. 2, Fig. S4). According to the sequence alignment, these residues correspond to Asn59, Arg73 and Val91, respectively, in TrxA, which contribute neutral or positive character to the equivalent surface region. Interestingly, Glu58 and Glu90 are present in all analyzed cyanobacterial TrxB proteins and represent distinctive features of this Trx type, whereas Glu72 is found in 75% of the analyzed cyanobacterial TrxB proteins (Mallén-Ponce et al., 2022). In TrxQ, a detailed comparison reveals an electropositive patch near the N-terminal region, mainly associated with the α-helix 1 and β-strand 2 (Fig. 2). This region contains several conserved basic residues, including Lys4, Lys5, Lys19, Arg48, Lys55, Lys60 and Lys80 (Fig. S4). Finally, TrxC is unique to cyanobacteria and contains a non-canonical active site (Fig. 2). Notably, it harbors a leucine residue instead of the conserved proline typically found between the two catalytic cysteines of canonical Trxs (López-Maury et al., 2018). This substitution likely alters the geometry of the active-site motif and may affect both structural stability and redox properties (Collet and Messens, 2010). In addition, structural modeling reveals a shorter and more distant β-strand 4 relative to the active site, associated with the absence of a conserved glycine residue corresponding to Gly92 in TrxA (Fig. 2). In TrxC, this position is occupied by a negatively charged Asp105, which interacts with the positively charged Arg107. Together with an additional Arg102-Glu104 interaction, these charged residues form a stabilizing network that may contribute to the distinctive folding of TrxC. These structural features may underlie the inability of TrxC to reduce substrates such as insulin or FBPase (López-Maury et al., 2018).

Three glycine residues are conserved among Trxs (Fig. S4). The first, located within the canonical CGPC active site motif, helps maintain the conformation of the catalytic site and contributes to determining the redox potential. The other two glycines flank β-strand 4 and define its length. These residues are structurally important because they occupy equivalent positions in turns found in other Trx structures (Collet and Messens, 2010).

Overall, this analysis indicates that *Synechocystis* Trxs share a conserved Trx fold but differ in surface properties that may contribute to functional diversification. TrxA and TrxB are the most structurally and electrostatically similar, although TrxB harbors specific acidic residues (Glu58, Glu72 and Glu90) that enhance the negative electrostatic potential surrounding the active site. These observations suggest that TrxB is the most suitable scaffold for dissecting the surface features that confer TrxA-specific function.

### Functional complementation of TrxA by chimeric thioredoxins

To investigate the determinants of TrxA-specific function, we constructed two *Synechocystis* mutant strains expressing TrxA/TrxB chimeric proteins. These hybrids were generated by interchanging the N- and C-terminal regions of TrxA and TrxB with respect to their active sites (Fig. 3a-b). Using the same strategy described above, the chimeric genes were placed under the control of the arsenite-inducible *arsB* promoter in a WT background, generating the STXABP and STXBAP strains (Fig. 3c). These resulting strains were verified by PCR analysis and shown to be fully segregated (Fig. 3d). Next, the endogenous *trxA* gene was disrupted in these strains by inserting a spectinomycin/streptomycin resistance cassette, generating the STXAB and STXBA strains (Fig. 3c). PCR analysis revealed incomplete segregation of the disrupted *trxA* locus in the STXAB strain, whereas the STXBA strain showed complete segregation (Fig. 3d). Similar segregation patterns were obtained when the same constructs were introduced into the SR background strain, further supporting the robustness of these findings (Fig. S6). These results indicate that the C-terminal region of TrxA contains determinants required for functional replacement of endogenous TrxA. Consistent with this finding, the C-terminal region contains several conserved motifs among cyanobacterial TrxA orthologs and chloroplast *m*-type Trxs (Mallén-Ponce et al., 2022).

**Figure 3.**
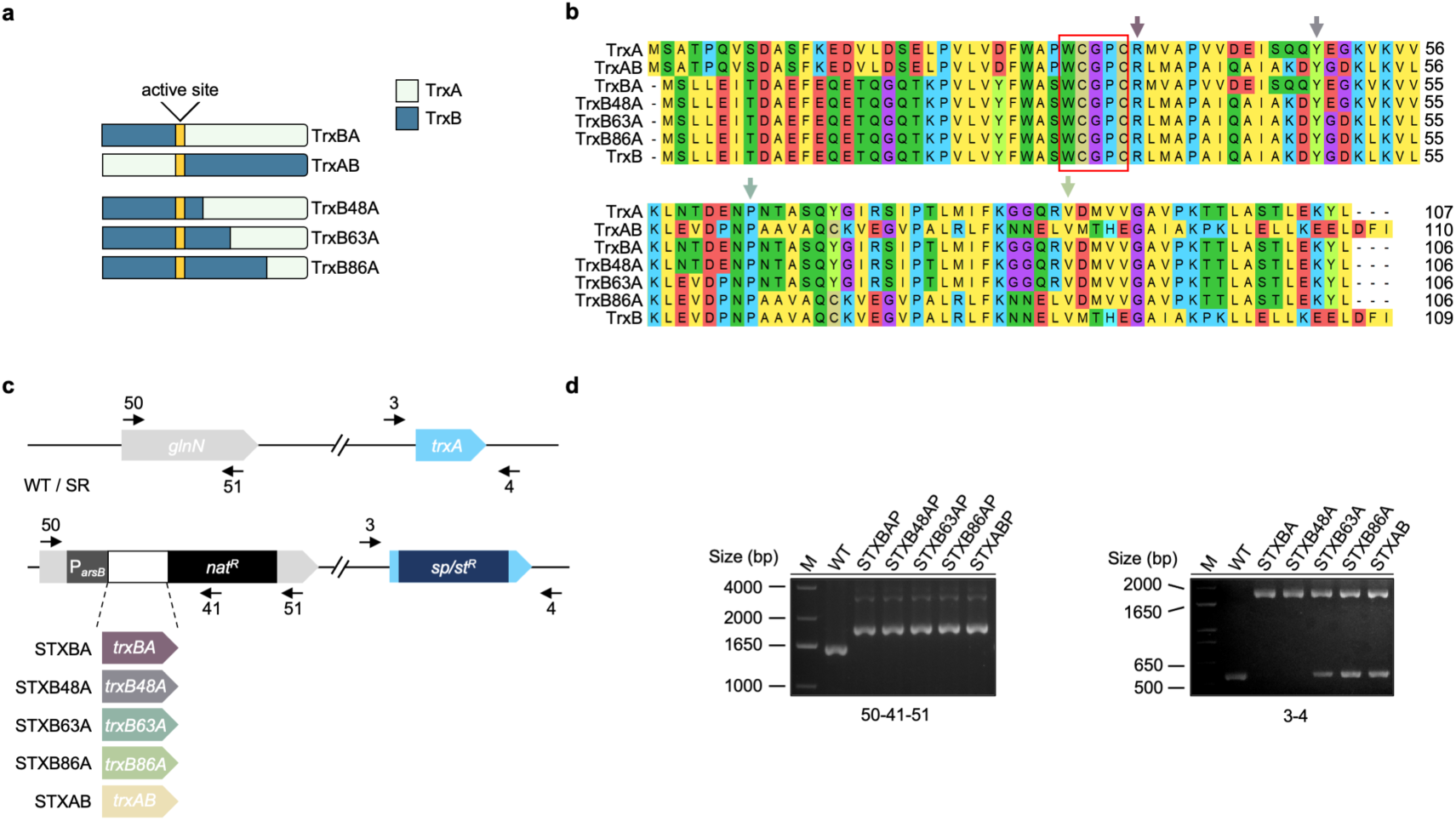
Construction and analysis of strains expressing TrxA/TrxB chimeric thioredoxins. **(a)** Schematic representation of TrxA, TrxB and the TrxA/TrxB chimeric Trxs. The catalytic active site and the regions derived from TrxA and TrxB are indicated. **(b)** Sequence alignment of TrxA, TrxB and the corresponding chimeric Trxs, showing the positions at which the TrxA and TrxB sequences were exchanged. Residue numbering and the active-site region are indicated. **(c)** Genetic strategy for the expression of the chimeric proteins under the control of the arsenite-inducible arsB promoter in the WT and SR genetic backgrounds. **(d)** PCR analysis of the parental strain and the corresponding derivatives carrying a disrupted *trxA* gene. Strains retaining the endogenous *trxA* gene are designated with the “P” suffix. M, molecular size marker. Alt text: A figure comprising several panel schematic diagrams of the *Synechocystis* chimeric strains, indicating the regions derived from each protein and the location of the active site, an amino acid sequence alignment highlighting the exchange sites, a diagram of the genetic strategy for expressing the chimeras using the arsB promoter and agarose gels showing the PCR products confirming the segregation of the strains.

To further explore the essential domains within the C-terminal region of TrxA, we generated additional mutant strains expressing TrxB/TrxA chimeric proteins. In these constructs, the N-terminal region of TrxB up to the active site was maintained, whereas increasing portions of the C-terminal region of TrxA were progressively replaced by the corresponding region of TrxB, generating STXB48AP, STXB63AP and STXB86AP strains (Figs. 3a-c). PCR analysis confirmed that these strains were fully segregated (Fig. 3d). Subsequently, the endogenous *trxA* gene was disrupted in these strains. PCR analysis revealed that, among these new chimeric strains, only STXB48A achieved complete segregation (Fig. 3d), suggesting that the residues essential for cell viability are not restricted to those located adjacent to the active site. A similar pattern was obtained when the SR strain was used as the genetic background (Fig. S6). The essential region retained in STXB48A (residues 49-106) corresponds to α-helices 3 and 4, as well as β-strands 3 and 4, and overlaps with highly conserved regions of TrxA (Mallén-Ponce et al., 2022). This contrasts with other Trxs, including TrxB, whose fully conserved region is restricted to a few amino acids near the active site (Mallén-Ponce et al., 2022).

To assess the functional impact of these structural differences, we examined the growth and phenotypic characteristics of the STXBA and STXB48A strains under induced and non-induced conditions. The WT and STXA3 strains were used as controls. Under induced conditions, there were no significant differences in growth between the STXBA strain and the control strains (Fig. 4a-b), whereas the STXB48A strain showed a slight growth delay. Interestingly, removal of the inducer caused a progressive growth retardation mainly in the STXBA strain (Fig. 4a-b), an effect not observed in the STXA3 strain. This difference may be related to the high initial intracellular levels of TrxA protein accumulated in the STXA3 strain under induced conditions (Fig. 1c). In addition, both STXBA and STXB48A strains displayed a distinct phenotypic change upon inducer removal, characterized by reduced levels of chlorophyll *a* and phycobilins (Fig. 4c, Fig. S7), whereas carotenoid levels were only slightly affected (Fig. 4c). These findings suggest that TrxA functionality is influenced by specific structural determinants located within the conserved C-terminal regions.

**Figure 4.**
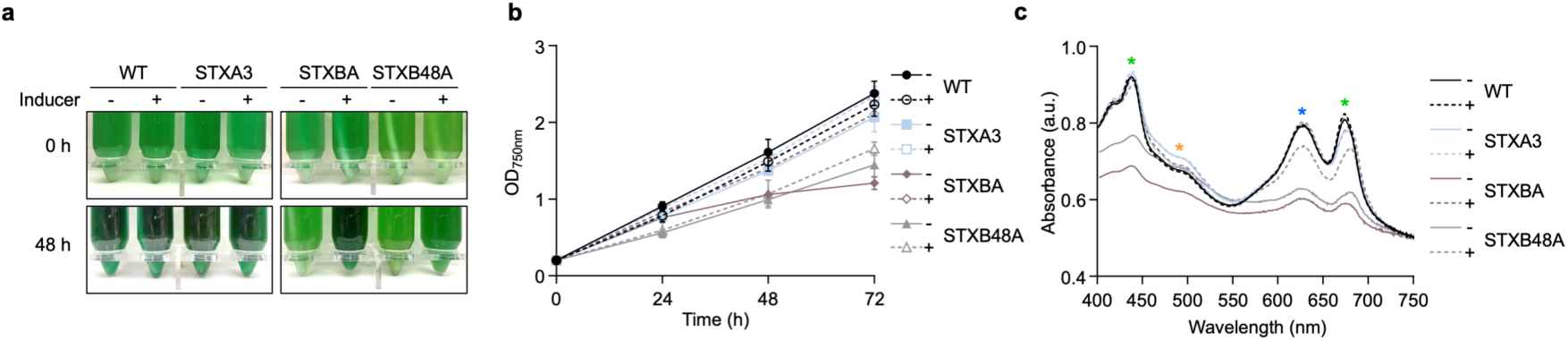
Growth and photosynthetic pigment profiles of strains expressing TrxA/TrxB chimeric thioredoxins. **(a)** Representative images of WT, STXA3, STXBA, and STXB48A cultures grown in the presence (+) or absence (−) of inducer for the indicated times. **(b)** Growth curves of the indicated strains under induced (+) and non-induced (−) conditions. **(c)** Whole-cell absorption spectra of WT, STXA3, STXBA and STXB48A strains grown after 48 h in the presence (+) or absence (−) of inducer. Asterisks indicate the absorption maxima corresponding to carotenoids, phycobiliproteins and chlorophyll. Alt text: Photographs of liquid cultures of WT, STXA3, STXBA and STXB48A strains grown with or without inducer over time, growth curves under both conditions, and whole-cell absorption spectra after 48 h with asterisks marking the carotenoid, phycobiliprotein and chlorophyll peaks.

### Characterization of the chimeric thioredoxins

The active site of Trxs is located in a surface-exposed loop at the beginning of α-helix 2, with the catalytic cysteine accessible to the solvent (Fig. 2). Residues near the active site contribute to Trx target recognition, including photosynthetic Trxs (Collet and Messens, 2010; Zaffagnini et al., 2018). To investigate the structural determinants underlying TrxA specificity, we analyzed the interaction of TrxA and the TrxA/TrxB chimeric proteins with the bifunctional enzyme FBP/SBPase from *Synechocystis*, a CB cycle enzyme essential under photoautotrophic conditions and previously identified as a TrxA target (Mallén-Ponce et al., 2021).

As a first step, we examined the molecular surfaces of the different chimeric proteins. *In silico* analysis predicted that TrxAB has a net charge of -8.1 and a theoretical pI of 4.7 (Table 1). Electrostatic surface modeling revealed an extended electronegative region surrounding the active site and α-helix 3, associated with the presence of acidic residues including Asp9, Glu59, Glu73 and Glu91, compared to TrxA and even TrxB (Fig. 5a). This distribution generates a continuous negative electrostatic potential around the active site. In addition, negatively charged residues in the N-terminal region of TrxAB, derived from the N-terminal region of TrxB, generate electronegative patches on the opposite side of the active site (Fig. 5a). In contrast to TrxAB, TrxBA has a predicted net charge and theoretical pI closer to those of TrxA (Table 1). Structural modeling revealed that the region surrounding the active site has a mixed polarity highly similar to TrxA (Fig. 5a). This similarity likely reflects the contribution of the TrxA derived C-terminal region, whereas TrxB N-terminal region is positioned away from the active site, where it has a more neutral character than the corresponding region in TrxA.

**Figure 5.**
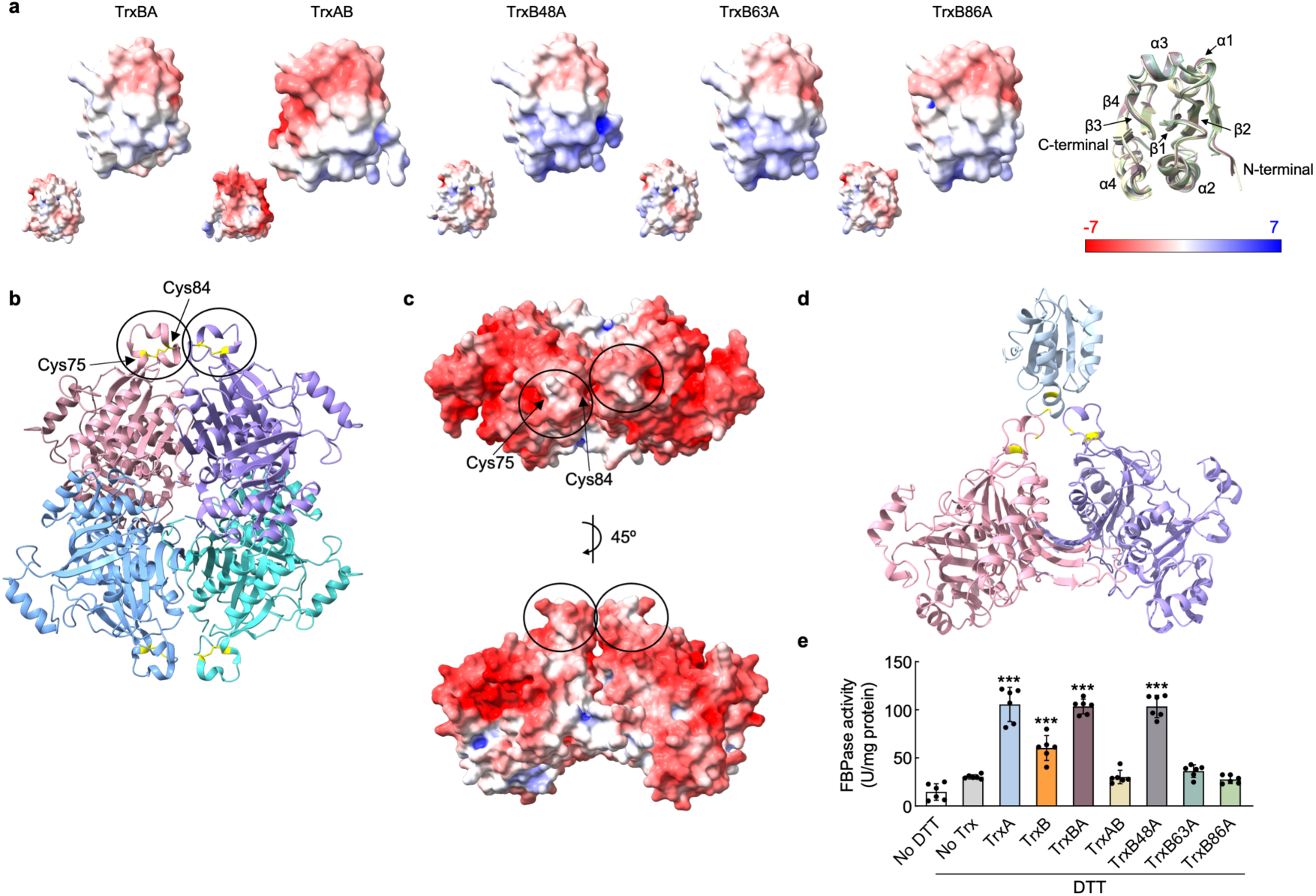
Structural and functional characterization of TrxA/TrxB chimeric thioredoxins. **(a)** Electrostatic surface representation of TrxA/TrxB chimeric Trxs predicted with AlphaFold, shown alongside a ribbon model highlighting secondary structural elements (α-helices and β-strands), as well as N-and C-terminus. Electrostatic potentials were calculated using the APBS algorithm and are shown from electropositive (blue) to electronegative (red). The smaller representations show the opposite face of each Trx. **(b)** Structural representation of the tetrameric FBP/SBPase, showing the four subunits and the flexible loop containing the regulatory cysteines Cys75 and Cys84. **(c)** Electrostatic surface representation of FBP/SBPase showing the region surrounding the regulatory cysteines Cys75 and Cys84, shown in two orientations (top view and 45° rotation). Electrostatic potentials were calculated using the APBS algorithm and are shown from electropositive (blue) to electronegative (red). **(d)** AlphaFold3-predicted model of the interaction between TrxA and dimeric FBP/SBPase, showing the proximity of the nucleophilic cysteine of TrxA to Cys84 of FBP/SBPase. **(e)** Trx-dependent activation of recombinant FBP/SBPase. Fructose-1,6-bisphosphatase activity was measured using untreated FBP/SBPase (−DTT) or after incubation with DTT alone (No Trx) or with DTT in the presence of TrxA, TrxB and the indicated TrxA/TrxB chimeric Trxs. Data points represent individual measurements, and error bars show the mean ± SD (*n* = 6). Asterisks represent significant differences relative to the No Trx control, as determined by one-way ANOVA followed by Dunnett’s multiple-comparisons test: ***P < 0.001; **P < 0.01; *P < 0.05; not significant, ≥0.05. Alt text: Molecular surface models of the TrxA/TrxB chimeric thioredoxins colored by electrostatic potential (blue positive, red negative) with ribbon diagrams, a structural model of the tetrameric FBP/SBPase highlighting the flexible loop with regulatory Cys75 and Cys84, its surface electrostatic potential around these cysteines in two orientations, and a predicted model of TrxA bound to dimeric FBP/SBPase. A dot plot that shows FBPase activity without DTT, with DTT alone, or with DTT plus TrxA, TrxB or each chimera, with asterisks marking significant differences from the DTT-only control.

We next analyzed TrxB48A, TrxB63A and TrxB86A, which exhibit less negative net charges and higher theoretical pI values than TrxA (Table 1). Structural modeling revealed that these chimeric Trxs retain the Trx fold but display marked differences in the electrostatic potential of their solvent-accessible surfaces compared with native TrxA (Fig. 2, Fig. 5a).

Sequence and structural analysis uncovered differences among the chimeric proteins that may contribute to TrxA function. In TrxB48A, α-helix 2 lacks the two negatively charged residues present in TrxA, Asp42 and Glu43 (Fig. 3b). Consistently, this region displays reduced negative electrostatic potential compared with TrxBA, which maintains a surface polarity similar to that of TrxA (Fig. 5a). However, these electrostatic differences do not appear to be essential for viability, as fully segregated strains lacking TrxA were obtained from cells expressing either TrxBA or TrxB48A.

Modelling TrxB63A revealed a more extended electronegative region around the active site (Fig. 5a). Because the region downstream of residue 63 is derived from TrxA, this electrostatic redistribution may reflect the combined contribution of acidic residues retained within the TrxB-derived N-terminal region, together with changes in their orientation and solvent exposure in the chimeric structure (Fig. 3b). Previous studies showed that substitutions of acidic residues within this region increased the efficiency of chloroplast FBPase activation, supporting a role for local electrostatic properties in target recognition (Eklund et al., 1991; de Lamotte-Guery et al., 1991; Wangensteen et al., 2001). The negative electrostatic character is further enhanced in TrxB86A, which retains the TrxB-derived Glu72 at the position equivalent to the positively charged Arg73 of TrxA (Fig. 3b, Fig. S4). This position is located within a region highly conserved among cyanobacterial TrxA orthologs and chloroplast *m*-type Trxs (Mallén-Ponce et al., 2022), and may contribute to the structural organization of the target-recognition surface. Overall, although all chimeric proteins retain the canonical Trx fold and active-site motif, notable differences are observed in the electrostatic potential of their solvent-accessible surfaces.

Several studies have suggested that protein-protein interactions between Trxs and their targets proteins depend on complementary electrostatic surfaces (Soulié et al., 1985; Rodriguez-Suarez et al., 1997; Mora-García et al., 1998; Collin et al., 2003; Balmer et al., 2004). We therefore examined the potential impact of changes in the electrostatic potential around the active site using the enzyme FBP/SBPase, a previously identified TrxA target in *Synechocystis* (Mallén-Ponce et al., 2021). Crystal structures of FBP/SBPase from *Synechocystis* and *Synechococcus* revealed the structural basis for its redox regulation, in which Cys75 and Cys84 form a regulatory disulfide bond (Feng et al., 2014; Cotton et al., 2015). However, biochemical studies in *Synechocystis* suggested that Cys75 and Cys99 may rearrange to form a disulfide bond in the inactive enzyme state (Feng et al., 2014). Consistently, *in vivo* redox analysis of FBP/SBPase showed higher oxidation levels in a TrxA knockdown mutant than in WT, resulting in reduced enzyme activity (Mallén-Ponce et al., 2021). Given the differences observed in the electrostatic properties of the chimeric Trxs (Fig. 5a), we used AlphaFold 3 to generate a computational model of the TrxA-FBP/SBPase interaction. The predicted models showed robust quality metrics (ipTM and pTM scores >0.7 and a favorable predicted aligned error (PAE) plot), supporting the reliability of the predicted interaction. As previously described, *Synechocystis* FBP/SBPase adopts a tetrameric structure formed by C1-C4 and C2-C3 dimer pairs, each composed of two monomers (Feng et al., 2014). The redox-regulated Cys75-Cys84 disulfide bond is located in a partially flexible loop between two β-sheets. In our model, the surface electrostatic potential calculated using the APBS algorithm revealed an extended electronegative surface around the regulatory Cys residues of FBP/SBPase (Fig. 5c).

We next performed molecular docking analysis between TrxA and dimeric FBP/SBPase to investigate their potential interactions. Alphafold 3 predicted several docking models of TrxA on the FBP/SBPase surface. In the four highest-scoring models, the nucleophilic Cys of the TrxA active site was positioned close to the regulatory Cys84 of FBP/SBPase (Fig. 5d). To further investigate the TrxA-FBP/SBPase interaction, we performed clustering analysis using PICKCLUSTER (Genz et al., 2023). This analysis identified potential interaction interfaces between TrxA and FBP/SBPase, highlighting residues involved in coordinating TrxA binding to the target protein. Across the predicted models, the interaction network was consistently anchored by residues surrounding the TrxA active site (Fig. S8). Recognition of FBP/SBPase was primarily mediated by Cys32 of TrxA together with adjacent active-site residues (Fig. S8). This interaction was further supported by a conserved surface region encompassing residues Gly71-Thr77, which includes Arg73 (Fig. S9). Several models also suggested additional contact regions involving charged residues such as Asp61 (Fig. S8). In all cases, these regions are highly conserved among different TrxA (*m*-type) proteins and are likely directly involved in the interaction with their target proteins. In FBP/SBPase, residues within the flexible recognition loop adjacent to Cys84 (Asp79-Pro86) appear to stabilize the loop conformation and contribute to the interaction with TrxA (Fig. S9).

Molecular docking analysis of TrxB and dimeric FBP/SBPase provided similar results, revealing an interaction network involving three regions that include the active site in the different predicted models (Fig. S8). Unlike TrxA, these regions of TrxB contain several negatively charged residues, including Glu72 and Glu90, which are predicted to interact with positively charged residues of FBP/SBPase, such as Arg50 and Lys81.

We next analyzed the potential interactions between the chimeric Trxs and FBP/SBPase. Molecular docking analysis predicted that TrxBA interacted with FBP/SBPase through regions similar to those identified for TrxA across the different models. In contrast, only one of the four highest-scoring TrxAB models positioned its active site close to Cys84 of FBP/SBPase, consistent with a potential redox-regulatory interaction (Fig. S8). The remaining chimeric Trxs also showed similar predicted interaction regions, with some specific differences among them. The main differences between TrxB48A and TrxB63A involved the turn connecting β-strand 2 and α-helix 3. Changes in this region were associated with a rearrangement of the predicted contacts of TrxB63A and also TrxB86A with FBP/SBPase. Specifically, in the models where TrxB63A and TrxB86A interacted through the loop containing Cys85, Glu58 was positioned close to Lys81 of FBP/SBpase, among other predicted contacts. Interestingly, previous studies in *E. coli* and plants have shown that substitutions of residues in this region affect the activity of TrxA (*m*-type) (de Lamotte-Guery et al., 1991; Wangensteen et al., 2001; Yokochi et al., 2019).

Given the essential role of FBP/SBPase under phototrophic conditions and its redox regulation by TrxA, we next examined the effects of different Trxs on FBPase activity (Fig. 5e). In the absence of both DTT and Trxs, FBP/SBPase displayed only basal activity, whereas addition of TrxA or TrxB increased FBPase activity, with TrxA showing a markedly stronger effect. The TrxBA chimera promoted activation to a level similar to that observed with TrxA, whereas TrxAB failed to activate FBP/SBPase. These results are consistent with the idea that preservation of key TrxA-like interface regions favors efficient functional regulation of FBP/SBPase. Interestingly, subsequent analysis of the TrxB48A, TrxB63A and TrxB86A chimeras showed that TrxB48A retained an activation capacity comparable to that of TrxA, whereas TrxB63A and TrxB86A failed to activate the enzyme. Together, these results indicate that specific structural determinants within TrxA are required for efficient activation of FBP/SBPase. Moreover, the predicted structural rearrangements between TrxB48A and TrxB63A may help explain the loss of FBP/SBPase activation observed in TrxB63A and TrxB86A.

TrxA (*m*-type) also interacts with and regulates multiple target proteins in photosynthetic eukaryotes, including several CB cycle enzymes. Therefore, we analyzed the activation of pea chloroplast fructose-1,6-bisphosphatase by TrxA, TrxB and the different chimeric Trxs. Pea FBPase is a homotetramer composed of approximately 40 kDa subunits, each of which contains a regulatory disulfide bond between the conserved Cys153 and Cys173 residues on its surface (Chiadmi et al., 1999). Reduction of this disulfide enables activation by chloroplast *f-*and *m-*type Trxs under light conditions (López-Jaramillo et al., 1997; Neira et al., 2024). Previous studies have shown that Pea FBPase activation is primarily driven by electrostatic interactions, with hydrophobic interactions playing a secondary role (Pla and Lopez-Gorge, 1981; Soulié et al., 1985; Crawford et al., 1986; Mora-García et al., 1998; Wangensteen et al., 2001). Consistent with this, Pea FBPase displays a large electronegative surface surrounding the two redox-sensitive cysteines, a feature conserved among plant orthologs (Marcus and Harrsch, 1990).

Molecular docking analysis identified a TrxA-Pea FBPase model in which the nucleophilic Cys of TrxA was positioned close to Cys153 of Pea FBPase (Fig. S10b). The predicted interface involved residues adjacent to the TrxA active site, as well as residues located between α-helix 3 and β-strand 3 (Arg73-Thr77). A second flexible region, located between β-strand 4 and α-helix 4 (Gly92-Pro95), was also positioned near the interface. Together, these regions positioned TrxA close to Cys153 of Pea FBPase and to adjacent surface regions that may contribute to the recognition of Pea FBPase (Fig. S10b).

Docking simulations performed with TrxB and the chimeric Trxs did not identify clearly productive interaction near Cys153. Only one TrxB48A model approached the regulatory site, although the predicted distance was not compatible with an efficient redox interaction. To examine whether the activity of the chimeras was target dependent, we also tested their ability to activate pea FBPase *in vitro*. The activation pattern of Pea FBPase was broadly consistent with that observed for the cyanobacterial FBP/SBPase. TrxA, TrxBA, and TrxB48A displayed similar activation levels, whereas TrxB showed somewhat lower activity. Progressive replacement of the C-terminal region of TrxA with the corresponding region of TrxB resulted in a progressive loss of activation capacity, which was markedly reduced with TrxB63A and no longer detectable with TrxB86A (Fig. S10c).

## DISCUSSION

The inability to eliminate the single *trxA* gene in *Synechocystis*, as well as all Trx-*m* isoforms present in plant chloroplasts strongly supports the idea that these proteins perform non-redundant functions. However, TrxA (or Trx-*m*) is also the most abundant Trx both in cyanobacteria and chloroplasts, respectively (Florencio et al., 2006; Okegawa and Motohashi, 2015), raising the possibility that its essentiality could be linked to its high intracellular abundance. Given that *Synechocystis* encodes several additional Trxs, these proteins might compensate for the loss of TrxA if expressed at comparable levels. To address this possibility, strains overexpressing different Trxs were generated and evaluated for their ability to functionally substitute for TrxA.

Our results demonstrate that overexpression of TrxB, TrxQ or TrxC failed to compensate for the loss of TrxA (Fig. 1a, Fig. 1b), indicating that TrxA essentiality cannot be explained solely by protein abundance. Overexpression restored TrxA protein levels to values comparable to those in the WT strain, whereas TrxB and TrxQ accumulated to approximately 7- and 35-fold higher levels, respectively (Fig. 1c). These increases are consistent with the lower endogenous abundance previously reported for these Trxs (Florencio et al., 2006; Baers et al., 2019; Wang et al., 2022; Spät et al., 2023), bringing their levels into a range comparable to that of TrxA in the WT strain. Nevertheless, neither TrxB nor TrxQ could functionally replace TrxA under these conditions. Together, these findings strongly support the existence of structural and/or functional properties specific to TrxA that cannot be compensated for by other Trxs, even when the latter accumulate to substantially higher levels.

Heterologous expression of NTRC, an alternative thiol-reducing system, did not produce any detectable phenotype in either the WT strain or STXA2, which has reduced TrxA levels (Fig. S3d-e). This finding is consistent with the idea that TrxA performs non-redundant functions that cannot be compensated for simply by introducing an alternative source of reducing power. Nevertheless, the involvement of other redox systems, including glutaredoxins (Sánchez-Riego et al., 2013), cannot be excluded.

The physicochemical basis of Trx specificity in photosynthetic organisms has been extensively investigated, particularly in cyanobacterial TrxA, its chloroplast *m*-type ortholog and chloroplast Trx-*f*. Comparative biochemical analyses in plants showed that trx-*m* and trx-*f* exhibit distinct target preferences, with Trx-*m* more efficiently activating NADP-malate dehydrogenase and Trx-*f* preferentially activating FBPase (Wolosiuk et al., 1979). Crystallographic analyses of spinach Trx-*f* and Trx-*m* revealed differences in surface charge distribution and loop conformations surrounding the WCGPC active site motif (Capitani et al., 2000), while three-dimensional modeling suggested that subtle variations in electrostatic surface potential contribute to preferential target recognition (Collin et al., 2003). Similarly, crystallographic analyses of chloroplast Trx-*f* isoforms from *Chlamydomonas reinhardtii* revealed a distinctive electropositive surface surrounding the active site that complements the electronegative surface of their target proteins (Lemaire et al., 2018).

Our structural analyses revealed that the cyanobacterial Trxs TrxA, TrxB and TrxQ share a high degree of structural similarity with chloroplast Trxs, both in their canonical fold and in the electrostatic properties of the region surrounding the active site (Collin et al., 2003; Lemaire et al., 2018; Table 1; Fig. 2). Nevertheless, these proteins displayed clear differences in surface charge distribution and loop conformation (Fig. 2), particularly in regions flanking the catalytic motif, suggesting distinct capacities to interact with specific redox partners. The coexistence of overall structural conservation and local divergence is consistent with the idea that the molecular determinants underlying Trx-substrate specificity were already established in the prokaryotic ancestors of plastids and subsequently retained during chloroplast evolution.

Although the structures of many Trx target proteins have been resolved, relatively few Trx– target complexes have been characterized structurally, limiting our understanding of the interaction interfaces and molecular contacts that determine target specificity. Previous modeling and comparative analyses in *Chlamydomonas reinhardtii* showed that Trx-*f* has an electropositive crown surrounding the active site and a flexible hinge region downstream of α-helix 1, both of which may contribute to target recognition (Collin et al., 2003; Lemaire et al., 2018). By contrast, Trx-*m* displays a mixed electrostatic surface around its catalytic cysteines, whereas Trx-*x* and Trx-*y* are characterized by predominantly neutral or electronegative surfaces. These observations are consistent with the structural models generated in this study and support the idea that both the distribution of surface charges and its spatial arrangement relative to the active site contribute to Trx target specificity.

Consistent with these observations, the structural models generated in this study indicate that TrxA has a more balanced electrostatic surface, whereas TrxB (Trx-*x* type) and TrxQ (Trx-*y* type) display more localized charge distributions. The spatial arrangement of these charges appears to influence target recognition: TrxB displays a conserved expanded electronegative surface surrounding the active site, which may be less favorable for productive interactions with CB cycle enzymes. By contrast, TrxQ exhibits a localized basic patch near its N-terminal region that could affect its recognition of complementary target surfaces (Collin et al., 2003; Fig. 2). Consistent with their established roles, TrxB (Trx-*x* type) and TrxQ (Trx-*y* type) have been primarily associated with antioxidant functions and have shown lower efficiency in regulating CB cycle enzymes. These differences support the idea that subtle structural and electrostatic variations contribute to the functional diversification of Trxs in photosynthetic organisms (Collin et al., 2003; Collin et al., 2004; Pérez-Pérez et al., 2009).

In our study, the generation of chimeric Trxs provided direct evidence that the C-terminal region of TrxA plays a central role in defining its essential function. The ability of TrxBA, but not TrxAB, to complement the loss of TrxA indicates that structural determinants required for productive interactions with target proteins are associated with the C-terminal region. This asymmetry suggests that the region downstream of the active site contributes to positioning the catalytic cysteines in a configuration compatible with efficient electron transfer. Importantly, this region encompasses α-helices 3 and 4 and adjacent β-strands, which were further delimited as essential in our progressive chimeric constructs. Alterations within this segment led to loss of function despite preservation of the overall Trx fold, indicating that global structural integrity alone is not sufficient to sustain activity. This interpretation is consistent with previous mutagenesis and biochemical studies showing that substitutions of residues surrounding the active site can affect Trx-dependent activation of target enzymes, such as FBPase and NADP-malate dehydrogenase, without disrupting the canonical fold. Studies of plant Trx-m proteins have likewise shown that sequence variations in conserved regions can affect FBPase activation (López-Jaramillo et al., 1997). In particular, several *in vitro* studies have introduced substitutions that neutralize or remove negatively charged residues in regions flanking the active site, resulting in significant changes in target enzyme activation efficiency. These modifications alter the electrostatic environment and the orientation of the interacting surfaces, potentially affecting target recognition and positioning rather than the catalytic chemistry of the active-site cysteines (de Lamotte-Guery et al., 1991; López-Jaramillo et al., 1997; Wangensteen et al., 2001; Yokochi et al., 2019). Indeed, removal of acidic residues or introduction of basic residues has even been reported to enhanced target enzyme activation, highlighting the role of local charge distribution in modulating Trx specificity and catalytic efficiency.

The *in vitro* activation assays support the idea that Trx specificity depends on localized structural determinants rather than on conservation of the canonical Trx fold alone. The activation patterns observed with both *Synechocystis* FBP/SBPase and Pea FBPase were broadly consistent with the *in vivo* complementation results (Fig. 4, Fig. 5, Fig. S8), suggesting that structural features retained in functional chimeras contribute to productive interactions with physiologically relevant targets. Notably, TrxA, TrxBA, and TrxB48A displayed similar activation levels towards Pea FBPase, whereas TrxB showed somewhat lower activity (Fig. S10c). Further replacement of the C-terminal region of TrxA progressively impaired activation, which was markedly reduced with TrxB63A and no longer detectable with TrxB86A (Fig. S10c). This pattern supports a contribution of the C-terminal region of TrxA to Pea FBPase activation, while indicating that the extent of this contribution depends on the structural context of the chimera. However, the differences between the in vivo complementation and in vitro activation assays indicate that the structural requirements for TrxA-dependent essential functions cannot be fully inferred from the activation of a single target enzyme.

Overall, our results extend previous observations from single-residue substitutions to larger structural modules, demonstrating that the C-terminal region of TrxA acts as a functional unit integrating key structural features required for target recognition and efficient electron transfer. This modular behavior is further supported by the ability of the TrxBA chimera to restore function, indicating that transplantation of this region into a TrxB scaffold is sufficient to confer substantial TrxA-like functionality.

### Author contributions

M.J.M.-P., M.J.H. and F.J.F. plan and designed the research; M.J.M.-P., J.M-L. and M.J.H. performed the research; M.J.M.-P. and M.J.H. analyze the data; M.J.M.-P. wrote the article and M.J.H. and F.J.F. contributed the final article version; M.J.M.-P. and M.J.H. supervised the work; M.J.M.-P. and F.J.F. funding acquisition.

## Acknowledgements

We thank Marian Sahrawy for providing the pea FBPase expression construct and Luis López-Maury for providing the *Synechocystis* FBP/SBPase expression construct.

## Supplementary Material

**Figure S1**. Schematic representation of the *arsB* promoter region used for thioredoxin expression.

**Figure S2**. Construction and characterization of thioredoxin-overexpressing strains in the SR background.

**Figure S3**. Construction and characterization of NTRC-expressing strains.

**Figure S4**. Multiple sequence alignment of *Synechocystis* thioredoxins.

**Figure S5**. Multiple sequence alignment of TrxA and *m*-type thioredoxins from photosynthetic organisms.

**Figure S6**. PCR Verification of strains expressing TrxA/TrxB chimeric thioredoxins in the SR background.

**Figure S7**. Chlorophyll concentration in strains expressing TrxA/TrxB chimeric thioredoxins.

**Figure S8**. AlphaFold analysis of residue interactions between TrxA, TrxB and TrxA/TrxB chimeric thioredoxins with *Synechocystis* FBP/SBPase.

**Figure S9**. AlphaFold model of the TrxA-SyFBP/SBPase interface.

**Figure S10**. Structural and functional analysis of Pea FBPase activation by TrxA, TrxB and chimeric thioredoxins.

**Table S1.** Sequence identity among *Synechocystis* thioredoxins.

**Table S2.** Synechocystis strains used in this study.

**Table S3.** Oligonucleotides used in this study.

**Table S4.** Antibodies used in this study.

**Table S5.** Plasmids used in this study.

## Funding

This work was supported by grant PID2022-138317NB-I00 to F.J.F., financed by MCIU/AEI/10.13039/501100011033 and by "FEDER, Una manera de hacer Europa", and by Junta de Andalucía (group BIO-0284); and by a Ramón y Cajal fellowship to M.J.M.-P. (RYC2024-051248-I), funded by the Spanish Ministry of Science, Innovation and Universities.

### Conflicts of Interest

The authors declare no conflict of interest.

### Data availability statement

All data are incorporated into the article or its online supplementary material.

## Notes

### Competing Interest Statement

The authors have declared no competing interest.

