## Supplemental Data 1 for "Structural determinants underlie the essentiality of the *m*-type thioredoxin TrxA in *Synechocystis*"

### Supplementary Material

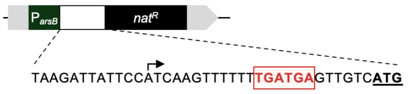


**Figure S1. Schematic representation of the *arsB* promoter region used for thioredoxin expression**. Nucleotide sequence and organization of the *arsB* promoter ($P_{arsB}$) indicating the conserved promoter regions and the original ribosome-binding site (RBS) retained upstream of the coding sequence.

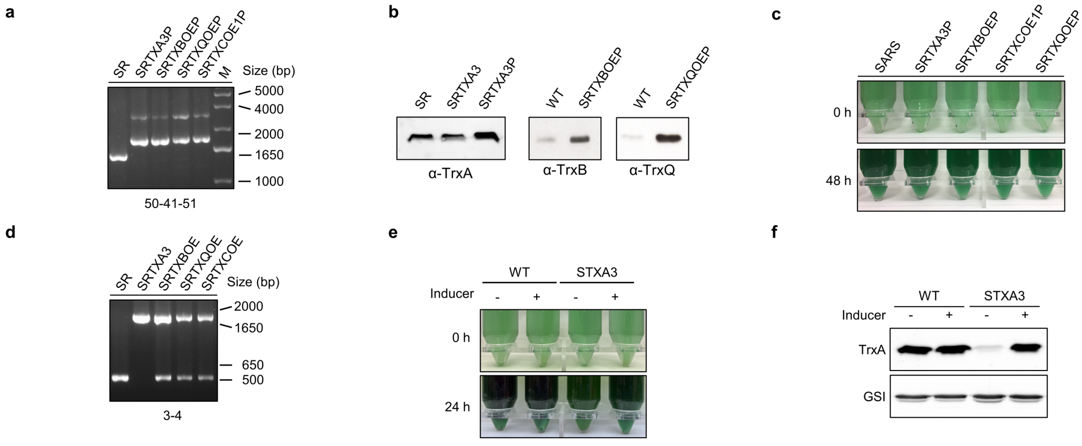


**Figure S2. Construction and characterization of thioredoxin-overexpressing strains in the SR background. (A)** PCR analysis of the SR-parental strain used to generate the corresponding trxA-disrupted derivatives. The numbers indicate the primers used. **(B)** Western blot analysis of TrxA, TrxB and TrxQ in the SR and trxA-disrupted strains. **(C)** Phenotype of WT, STXA3P, STXBOEP, STXCOE1P and STXQOEP strains at 0h and 48 h. **(D)** PCR analysis of *trxA* segregation in the indicated strains. M, molecular size marker. Strains carrying the disrupted *trxA* gene are named without the “P” suffix. **(E)** Representative images of WT and STXA3 cultures grown in the presence (+) or absence (−) of the inducer for the indicated times. **(F)** Western blot analysis of TrxA levels in WT and STXA3 cultures under induced (+) and non-induced (−) conditions.

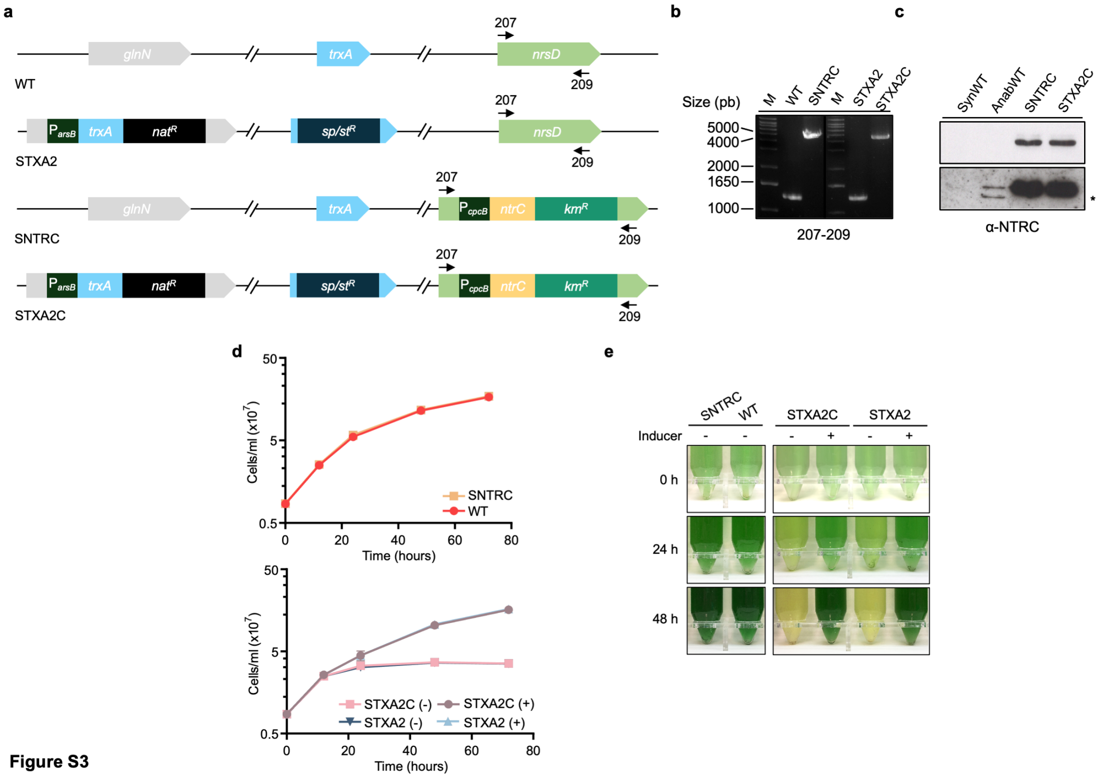


Figure S3

**Figure S3. Construction and characterization of NTRC-expressing strains. (A)** Genetic strategy for the expression of the *Anabaena* sp. PCC 7120 *ntrC* gene under the control of the constitutive *cpcB* promoter in the WT and STXA2 backgrounds. **(B)** PCR analysis confirming complete chromosome segregation of the **SNTRC** and **STXA2C** strains. **(C)** Western blot analysis of NTRC levels in the indicated strains. Western blots are representative of three independent biological replicates. **(D)** Growth curves of WT, SNTRC, STXA2 and STXA2C strains. **(E)** Representative images of WT, SNTRC, STXA2 and STXA2C strains at the indicated times. For panels (D) and (E), (+) and (−) indicate cultures grown in the presence or absence of arsenite, respectively.

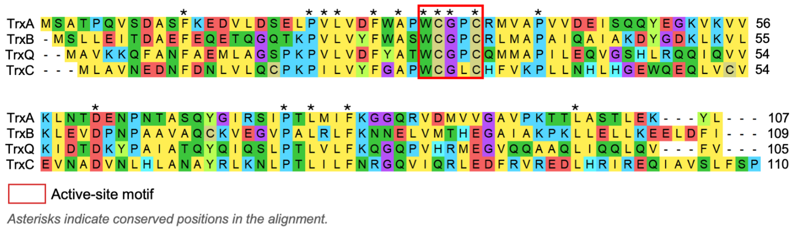


**Figure S4. Multiple sequence alignment of *Synechocystis* thioredoxins.** Multiple sequence alignment of TrxA, TrxB, TrxQ and TrxC from *Synechocystis*. Conserved residues are indicated by asterisks, and the active-site motifs are marked.

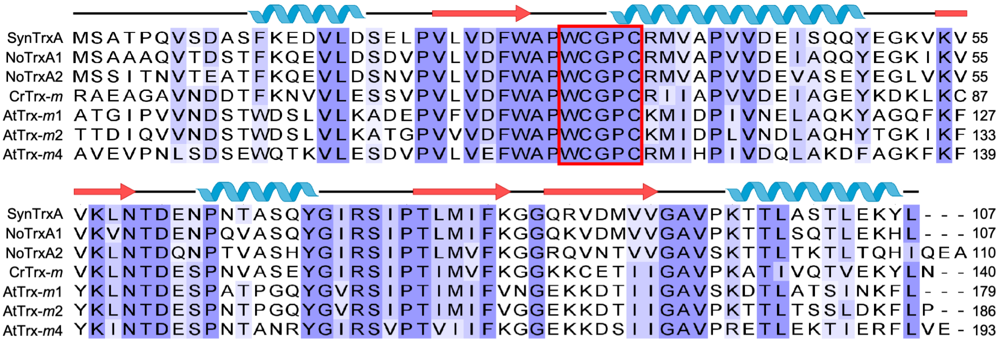

**Figure S5. Multiple sequence alignment of TrxA and *m*-type thioredoxins from photosynthetic organisms**. Multiple sequence alignment of *Synechocystis* sp. PCC 6803 TrxA (SynTrxA) and representative *m*-type thioredoxins from *Arabidopsis thaliana* (AtTrx-m1, AtTrx-m2 and AtTrx-m4), *Chlamydomonas reinhardtii* (CrTrx-m) and *Nostoc* sp. PCC 7120 (NoTrxA1 and NoTrxA2). The active-site motif and conserved regions located downstream of the active site are indicated with asterisks.

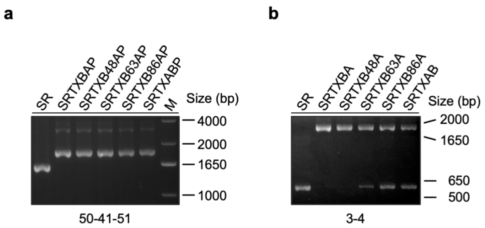


**Figure S6**. **PCR Verification of strains expressing TrxA/TrxB chimeric thioredoxins in the SR background.** (A) PCR analysis of the indicated SR-derived strains. The numbers indicate the primers used. (B) PCR analysis of *trxA* segregation in the indicated strains. M, molecular size marker. Strains carrying the disrupted *trxA* gene are named without the "P" suffix.

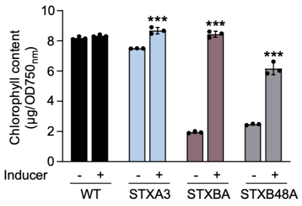


**Figure S7. Chlorophyll concentration in strains expressing TrxA/TrxB chimeric thioredoxins.** Chlorophyll content was measured in WT, STXA3, STXBA, and STXB48A cultures after 48 h in the presence (+) or absence (−) of inducer. Data points represent individual measurements, and error bars show the mean ± SD ($n$ = 3).

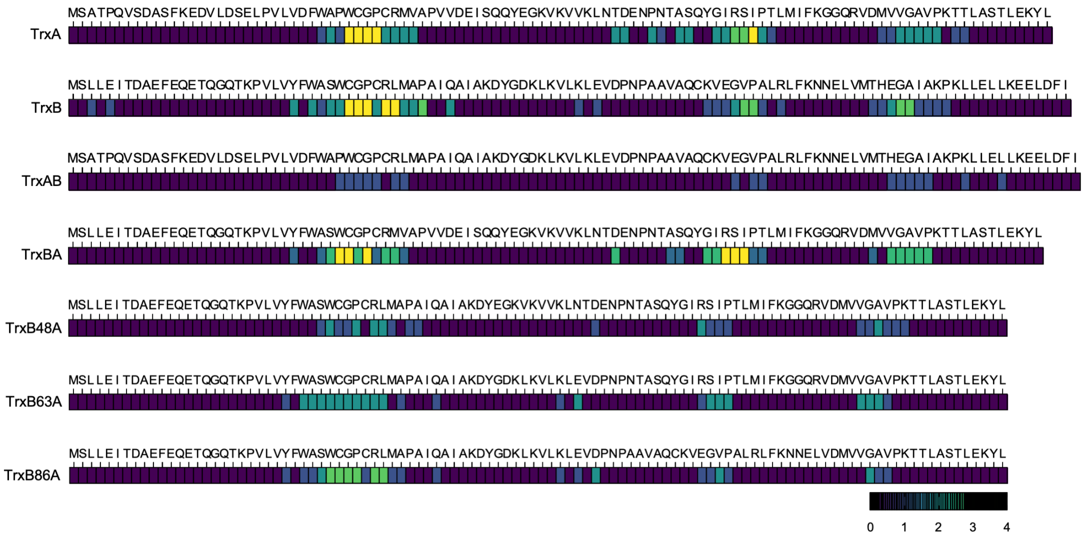


**Figure S8**. **AlphaFold analysis of residue interactions between TrxA, TrxB and TrxA/TrxB chimeric thioredoxins with FBP/SBPase.** Identification of interacting residues mapped across the models generated by AlphaFold and clustered using PICKLUSTER. The color scale indicates the number of models (from 1 to 4) in which each residue is predicted to participate in the protein-protein interaction.

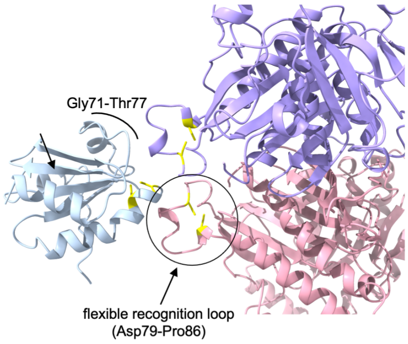


**Figure S9**. **AlphaFold model of the TrxA-FBP/SBPase interface**. Structural detail of the predicted complex obtained by AlphaFold, highlighting the Gly71–Thr77 region, the flexible recognition loop (Asp79–Pro86), and key interface residues (yellow). Interacting residues and catalytic/redox-active cysteines are highlighted in yellow.

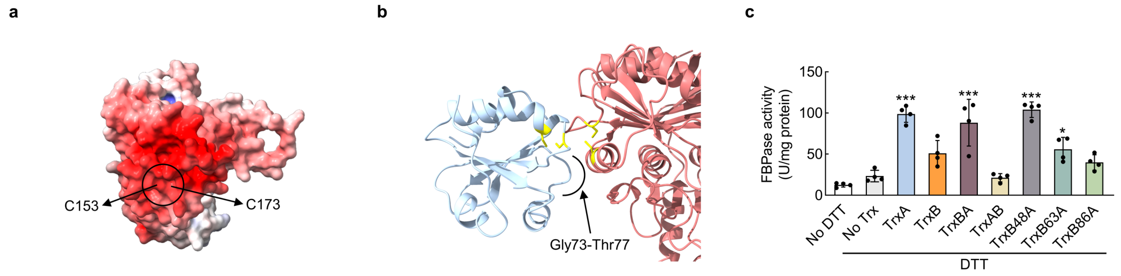


**Figure S10**. Structural and functional analysis of PsFBPase activation by TrxA, TrxB and chimeric thioredoxins. **(A)** Electrostatic surface representation of the AlphaFold predicted structure of PsFBPase showing the region surrounding the regulatory cysteines (C153 and C173). Electrostatic potentials were calculated using the APBS algorithm and are shown from electropositive (blue) to electronegative (red). **(B)** Detailed structural AlphaFold model of the predicted interaction interface between TrxA and PsFBP/SBPase obtained by AlphaFold and PICKLUSTER, highlighting the Gly73–Thr77 region and key participating residues (yellow). **(C)** Trx-dependent activation of recombinant PsFBP/SBPase. Fructose-1,6-bisphosphatase activity was measured using untreated PsFBP/SBPase (−DTT) or after incubation with DTT alone (No Trx) or with DTT in the presence of TrxA, TrxB and the indicated TrxA/TrxB chimeric Trxs. Data points represent individual measurements, and error bars show the mean ± SD ($n = 4$). Asterisks represent significant differences relative to the No Trx control, as determined by one-way ANOVA followed by Dunnett's multiple-comparisons test: ***$P < 0.001$; **$P < 0.01$; *$P < 0.05$; not significant, $\geq 0.05$.

**Table S1**. Sequence identity (%) among *Synechocystis* Trxs. Values correspond to the pairwise sequence comparisons shown in Figure S4.

| | **TrxA** | **TrxB** | **TrxQ** | **TrxC** |
|---|---|---|---|---|
| **TrxA** | — | 37.74 | 39.05 | 28.85 |
| **TrxB** | | — | 32.38 | 21.50 |
| **TrxQ** | | | — | 29.81 |
| **TrxC** | | | | — |

**Table S2**. *Synechocystis* strains used in this study

| STRAIN | BACKGROUND | GENOTYPE | RESISTANCE | SOURCE |
|---|---|---|---|---|
| WT | Synechocystis sp. PCC 6803 | Wild type | — | Laboratory collection |
| SR (ΔarsR; SARSR) | Synechocystis sp. PCC 6803 | ΔarsR | CmR | López-Maury et al., 2003 |
| STXA3P | WT | glnN::ParsB:trxA:NatR | NatR | This study |
| STXBOEP | WT | glnN::ParsB:trxB:NatR | NatR | This study |
| STXQOEP | WT | glnN::ParsB:trxQ:NatR | NatR | This study |
| STXCOE1P | WT | glnN::ParsB:trxC:NatR | NatR | This study |
| SRTXA3P | SR | ΔarsR::Cm, glnN::ParsB:trxA:NatR | NatR, CmR | This study |
| SRTXBOEP | SR | ΔarsR::Cm, glnN::ParsB:trxB:NatR | NatR, CmR | This study |
| SRTXQOEP | SR | ΔarsR::Cm, glnN::ParsB:trxQ:NatR | NatR, CmR | This study |
| SRTXCOE1P | SR | ΔarsR::Cm, glnN::ParsB:trxC:NatR | NatR, CmR | This study |
| STXA3 | STXA3P | glnN::ParsB:trxA:NatR; ΔtrxA::sp/stR | NatR, Sp/StR | This study |
| STXBOE | STXBOEP | glnN::ParsB:trxB:NatR; ΔtrxA::sp/stR | NatR, Sp/StR | This study |
| STXQOE | STXQOEP | glnN::ParsB:trxQ:NatR; ΔtrxA::sp/stR | NatR, Sp/StR | This study |
| STXCOE1 | STXCOE1P | glnN::ParsB:trxC:NatR; ΔtrxA::sp/stR | NatR, Sp/StR | This study |
| SRTXA3 | SRTXA3P | ΔarsR::Cm, glnN::ParsB:trxA:NatR; ΔtrxA::sp/stR | NatR, CmR, Sp/StR | This study |
| SRTXBOE | SRTXBOEP | ΔarsR::Cm, glnN::ParsB:trxB:NatR; ΔtrxA::sp/stR | NatR, CmR, Sp/StR | This study |
| SRTXQOE | SRTXQOEP | ΔarsR::Cm, glnN::ParsB:trxQ:NatR; ΔtrxA::sp/stR | NatR, CmR, Sp/StR | This study |
| SRTXCOE1 | SRTXCOE1P | ΔarsR::Cm, glnN::ParsB:trxC:NatR; ΔtrxA::sp/stR | NatR, CmR, Sp/StR | This study |
| STXABP | WT | glnN::ParsB:trxAB:NatR | NatR | This study |
| STXBAP | WT | glnN::ParsB:trxBA:NatR | NatR | This study |
| STXB48AP | WT | glnN::ParsB:trxB48A:NatR | NatR | This study |
| STXB63AP | WT | glnN::ParsB:trxB63A:NatR | NatR | This study |
| STXB86AP | WT | glnN::ParsB:trxB86A:NatR | NatR | This study |
| STXAB | STXABP | glnN::ParsB:trxAB:NatR; ΔtrxA::sp/stR | NatR, Sp/StR | This study |
| STXBA | STXBAP | glnN::ParsB:trxBA:NatR; ΔtrxA::sp/stR | NatR, Sp/StR | This study |
| STXB48A | STXB48AP | glnN::ParsB:trxB48A:NatR; ΔtrxA::sp/stR | NatR, Sp/StR | This study |
| STXB63A | STXB63AP | glnN::ParsB:trxB63A:NatR; ΔtrxA::sp/stR | NatR, Sp/StR | This study |
| STXB86A | STXB86AP | glnN::ParsB:trxB86A:NatR; ΔtrxA::sp/stR | NatR, Sp/StR | This study |
| SRTXABP | SR | ΔarsR::Cm, glnN::ParsB:trxAB:NatR | NatR, CmR | This study |
| SRTXBAP | SR | ΔarsR::Cm, glnN::ParsB:trxBA:NatR | NatR, CmR | This study |
| SRTXB48AP | SR | ΔarsR::Cm, glnN::ParsB:trxB48A:NatR | NatR, CmR | This study |
| SRTXB63AP | SR | ΔarsR::Cm, glnN::ParsB:trxB63A:NatR | NatR, CmR | This study |
| SRTXB86AP | SR | ΔarsR::Cm, glnN::ParsB:trxB86A:NatR | NatR, CmR | This study |
| SRTXAB | SRTXABP | ΔarsR::Cm, glnN::ParsB:trxAB:NatR; ΔtrxA::sp/stR | NatR, CmR, Sp/StR | This study |
| SRTXBA | SRTXBAP | ΔarsR::Cm, glnN::ParsB:trxBA:NatR; ΔtrxA::sp/stR | NatR, CmR, Sp/StR | This study |
| SRTXB48A | SRTXB48AP | ΔarsR::Cm, glnN::ParsB:trxB48A:NatR; ΔtrxA::sp/stR | NatR, CmR, Sp/StR | This study |
| SRTXB63A | SRTXB63AP | ΔarsR::Cm, glnN::ParsB:trxB63A:NatR; ΔtrxA::sp/stR | NatR, CmR, Sp/StR | This study |
| SRTXB86A | SRTXB86AP | ΔarsR::Cm, glnN::ParsB:trxB86A:NatR; ΔtrxA::sp/stR | NatR, CmR, Sp/StR | This study |
| SNTRC | WT | nrsD::PcpcB:ntrC:kmR | KmR | This study |
| STXA2 | WT | glnN::ParsB:trxA:NatR; ΔtrxA::sp/stR | NatR, Sp/StR | Mallén-Ponce et al., 2021 |
| STXA2C | STXA2 | glnN::ParsB:trxA:NatR; nrsD::PcpcB:ntrC:kmR; ΔtrxA::sp/stR | NatR, KmR, Sp/StR | This study |
| AnabWT | Anabaena sp. PCC 7120 | Wild type | — | Laboratory collection |

**Table S3.** Oligonucleotides used in this study

| Oligonucleotide | Sequence (5′–3′) |
|---|---|
| GlnN_check_F (50) | ATGCAGGCCAGTCTTCCTAA |
| GlnN_check_R (51) | AAATGGCAGTGTCCAAGTCC |
| Prom_Ck1_R (41) | AAGAGTGGTACCCATGGTAAACGATCCTCATCCTG |
| TrxA_1(3) | TAGGAGCGCCATATGAGTGCTACCCCTCAAGTTTCC |
| TrxA_2(4) | AAACCCCGGGAAGTGATTGGCGAATTAG |
| NrsD_BssHII_R (207) | AAGCGCGCCTCGGAGTCATGGTGATGGG |
| NrsD_BssHII_F (209) | AAGCGCGCCTTTCACTGCTTGCGGAACC |
| ParsB_EcoRI_F | TTCGAATTCTCGTTTCTTGGCTCATACC |
| PArsB_TrxA_5'_R | GTAGCACTCATGACAACTCATCAAAAAAACTTGAT |
| PArsB_TrxA_5'_F | GATGAGTTGTCATGAGTGCTACCCCTCAAGTTTCC |
| PArsB_TrxB_5'_F | GATGAGTTGTCATGAGTTTACTGGAAATC |
| PArsB_TrxB_5'_R | GTAAACTCATGACAACTCATCAAAAAAACTTGATG |
| PArsB_TrxC_5'_F | GATGAGTTGTCATGGTTGTAGCATGGACAAC |
| PArsB_TrxC_5'_R | GCTACAACCATGACAACTCATCAAAAAAACTTGAT |
| PArsB_TrxQ_5'_F | GATGAGTTGTCATGGCCGTCAAAAAACAGTTTG |
| PArsB_TrxQ_5'_R | TTGACGGCCATGACAACTCATCAAAAAAACTTGAT |
| TrxB_NotI | AATTGCGGCCGCAGAAGGATTATAAAC |
| TrxB_NdeI | AATTCATATGAGTTTACTGGAAATCAC |
| TrxA_NdeI | AATTCATATGAGTGCTACCCCTCAAG |
| TrxA_NotI | AATTGCGGCCGCCACAAAGCAGGGAAGCCAG |
| TrxC_NotI_R | AATTAATTGCGGCCGCTTATCAGGGAGAGAAAAG |
| TrxQ_NotI_R | AATTAATTGCGGCCGCTTATTACACAAAAAACCTG |
| AC_TrxA_F | GGTGTGGCCCTTGTCGCATGG |
| AC_TrxA_R | GCGACAAGGGCCACACCAGGAG |
| AC_TrxB_F | GGTGTGGCCCTTGTCGGTTGATG |
| AC_TrxB_R | CCGACAAGGGCCACACCAGGGAG |
| TrxB_3´_R | AATTAATTGCGGCCGCTTATTAAATAAAATCCAATT |
| TrxA_3´_R | AATTAATTGCGGCCGCTTATTAAAGATATTTTTCTA |
| TrxB144_TrxA148_F | GCCAAGGATTATGAAGGTAAGGTCAAAGTGG |
| TrxB144_TrxA148_R | GACCTTACCTTCATAATCCTTGGCGATCGCCCTGG |
| TrxB189_TrxA193_F | GATCCCAACCCTAATACCGCTAGTCAGTATG |
| TrxB189_TrxA193_R | ACTAGCGGTATTAGGGTTGGGATCCACTTC |
| TrxB258_TrxA262_F | AATGAGCTAGTGGATATGGTGGTGGGAGCCG |
| TrxB258_TrxA262_R | CACCACCATATCCACTAGCTCATTATTTTGAAC |

**Table S4.** Antibodies

| Antibody | Target protein | Dilution |
|---|---|---|
| α-TrxA | TrxA | 1:3000 |
| α-TrxB | TrxB | 1:2000 |
| α-TrxQ | TrxQ | 1:2000 |
| α-NTRC | NTRC | 1:1000 |
| α-GSI | Glutamine synthetase I (GSI) | 1:250000 |

### **Table S5.** Plasmids used in this study

| Plasmid | Description / relevant features | Antibiotic resistance | Source / reference |
|---|---|---|---|
| pET28a(+) | E. coli expression vector; N-terminal His-tag; T7 promoter | Kanamycin | Novagen |
| pglnN::ParsB:natR | pBS-based vector; glnN (slr0288) locus disrupted by the arsenite-inducible arsB promoter and a nourseothricin resistance cassette | Ampicillin ; nourseothricin | [Mallén-Ponce et al., 2022] |
| pnrsD::PcpcB:Km | pBS-based vector; nrsD (slr0796) locus disrupted by the constitutive cpcB promoter and a kanamycin resistance cassette | Ampicillin ; nourseothricin | Kindly provided by L. López-Maury |
| pGQ5.4+ | Disrupts the endogenous trxA gene with a spectinomycin/streptomycin resistance cassette | Spectinomycin/streptomycin | Mallén-Ponce et al., 2022 |
| pglnN::ParsB-trxA:natR | trxA under the control of ParsB, retaining the original RBS | Nourseothricin | This study |
| pglnN::ParsB-trxB:natR | trxB under the control of ParsB | Nourseothricin | This study |
| pglnN::ParsB-trxQ:natR | trxQ under the control of ParsB | Nourseothricin | This study |
| pglnN::ParsB-trxC:natR | trxC under the control of ParsB | Nourseothricin | This study |
| pglnN::ParsB-trxAB:natR | TrxA/TrxB chimera (TrxA N-terminus, TrxB C-terminus) under ParsB | Nourseothricin | This study |
| pglnN::ParsB-trxBA:natR | TrxB/TrxA chimera (TrxB N-terminus, TrxA C-terminus) under ParsB | Nourseothricin | This study |
| pglnN::ParsB-trxB48A:natR | TrxB residues 1–48 + TrxA residues 50–end under ParsB | Nourseothricin | This study |
| pglnN::ParsB-trxB63A:natR | TrxB residues 1–63 + TrxA residues 65–end under ParsB | Nourseothricin | This study |
| pglnN::ParsB-trxB86A:natR | TrxB residues 1–86 + TrxA residues 88–end under ParsB | Nourseothricin | This study |
| pET28_trxA | TrxA for recombinant production (N-terminal His-tag) | Kanamycin | Pérez-Pérez et al. |
| pET28_trxB | TrxB for recombinant production (N-terminal His-tag) | Kanamycin | Pérez-Pérez et al. |
| pET28_trxAB | TrxAB chimera for recombinant production | Kanamycin | This study |
| pET28_trxBA | TrxBA chimera for recombinant production | Kanamycin | This study |
| pET28_trxB48A | TrxB48A chimera for recombinant production | Kanamycin | This study |
| pET28_trxB63A | TrxB63A chimera for recombinant production | Kanamycin | This study |
| pET28_trxB86A | TrxB86A chimera for recombinant production | Kanamycin | This study |
| pET28-FBPase | Pea chloroplast FBPase for recombinant production | Kanamycin | Kindly provided by M. Sahrawy |
| pET28-FBP/SBPase | Synechocystis FBP/SBPase for recombinant production | Kanamycin | Kindly provided by L. López-Maury |
| p28AnNTR2 | pET28 derivative carrying Anabaena sp. PCC 7120 ntrC (all0737) | Kanamycin | lab |
| pBS-NTRC | ntrC (all0737) under the constitutive cpcB promoter in the nrsD locus | Ampicillin; kanamycin | This study |
